## Supplementary information for "Longitudinal transcriptional changes reveal genes from the natural killer cell-mediated cytotoxicity pathway as critical players underlying COVID-19 progression"

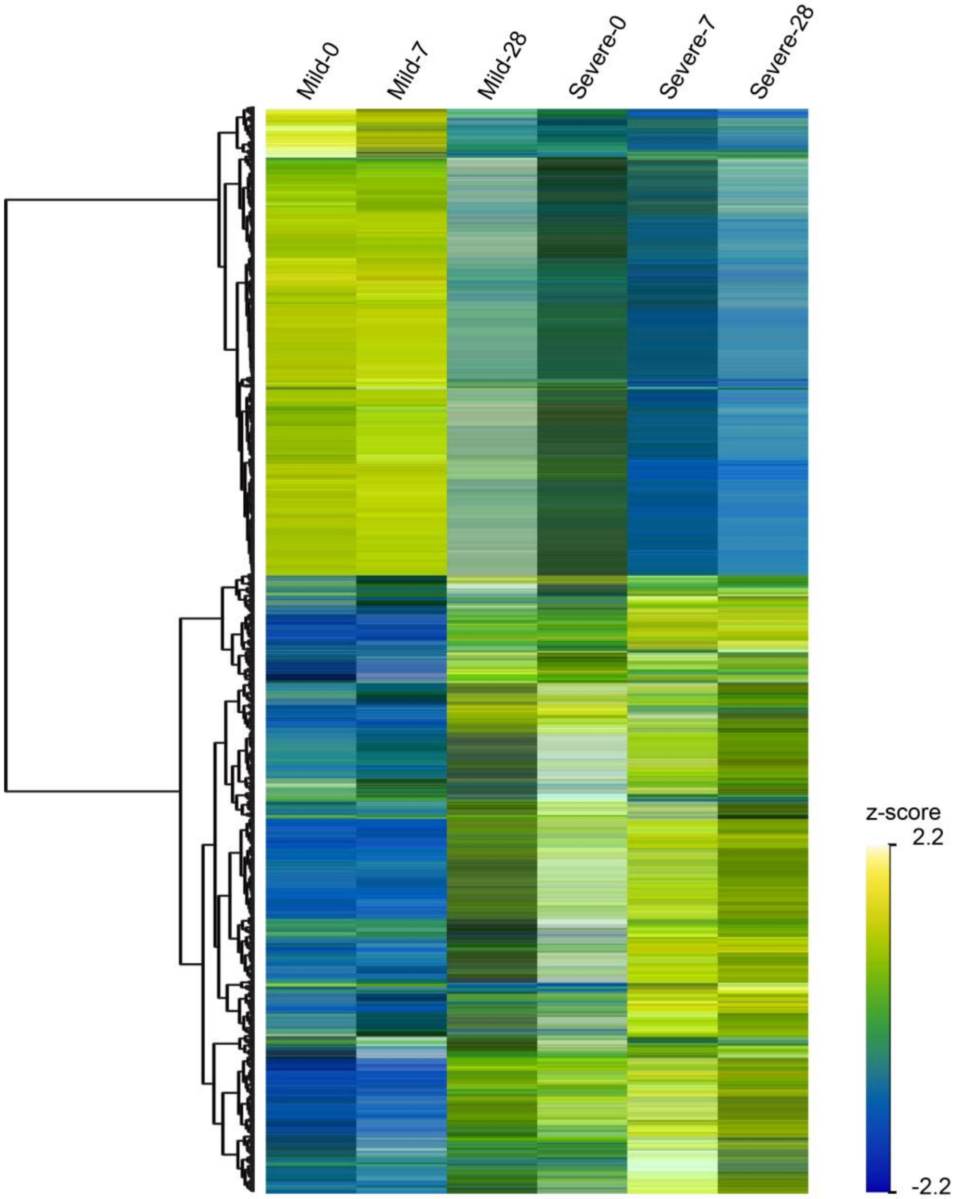


**Figure 1 - figure supplement 1.** Heatmap of temporally and differentially expressed genes over the course of COVID-19 progression. At the top, each column corresponds to the sampling points (0, 7, and 28 days since recruitment) of mild and severe patients. Genes are displayed as horizontal rows and are clustered by the similarity of expression profiles, represented by the dendrogram to the left of the heatmap. To the right of the heatmap, yellowish color indicates higher expression, while bluish color means lower expression represented by the z-score of normalized read counts.

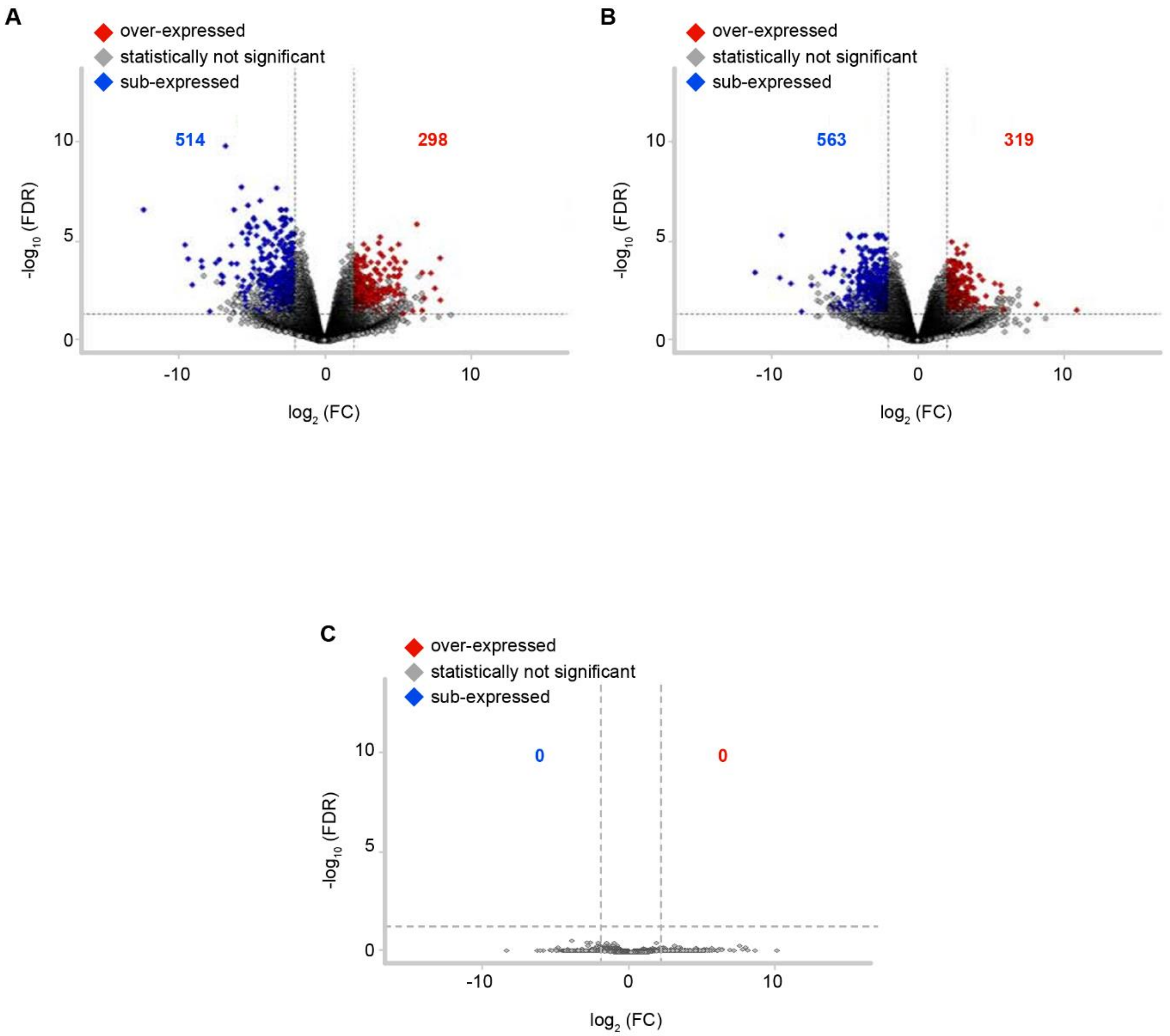


**Figure 1 - figure supplement 2.** Volcano plot depicting pairwise gene expression comparisons for detected DEGs between mild and severe COVID-19 patients at D0, D7 and D28 (**A**-**B**-**C**, respectively). Red or blue indicate genes that were significantly over- or sub-expressed at a particular sampling time, based on filtering by FDR ≤ 0.05 and the absolute value of -log10 (FC) (≥ 2.0). The remaining genes that do not show differential expression are indicated in gray. FDR = False discovery rate; FC = Fold change.

23

**A**

KEGG graph of Th1 and Th2 cell differentiation

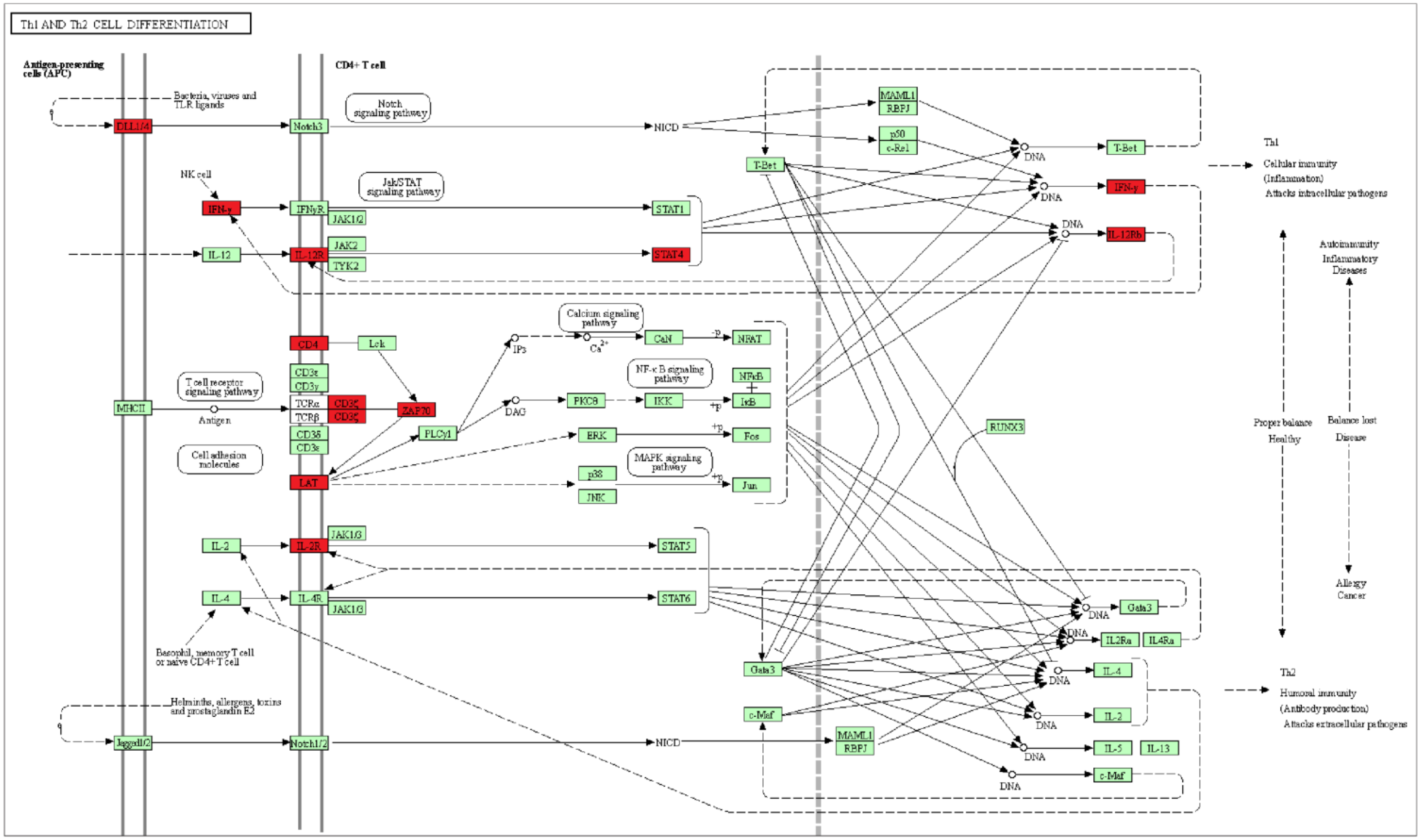

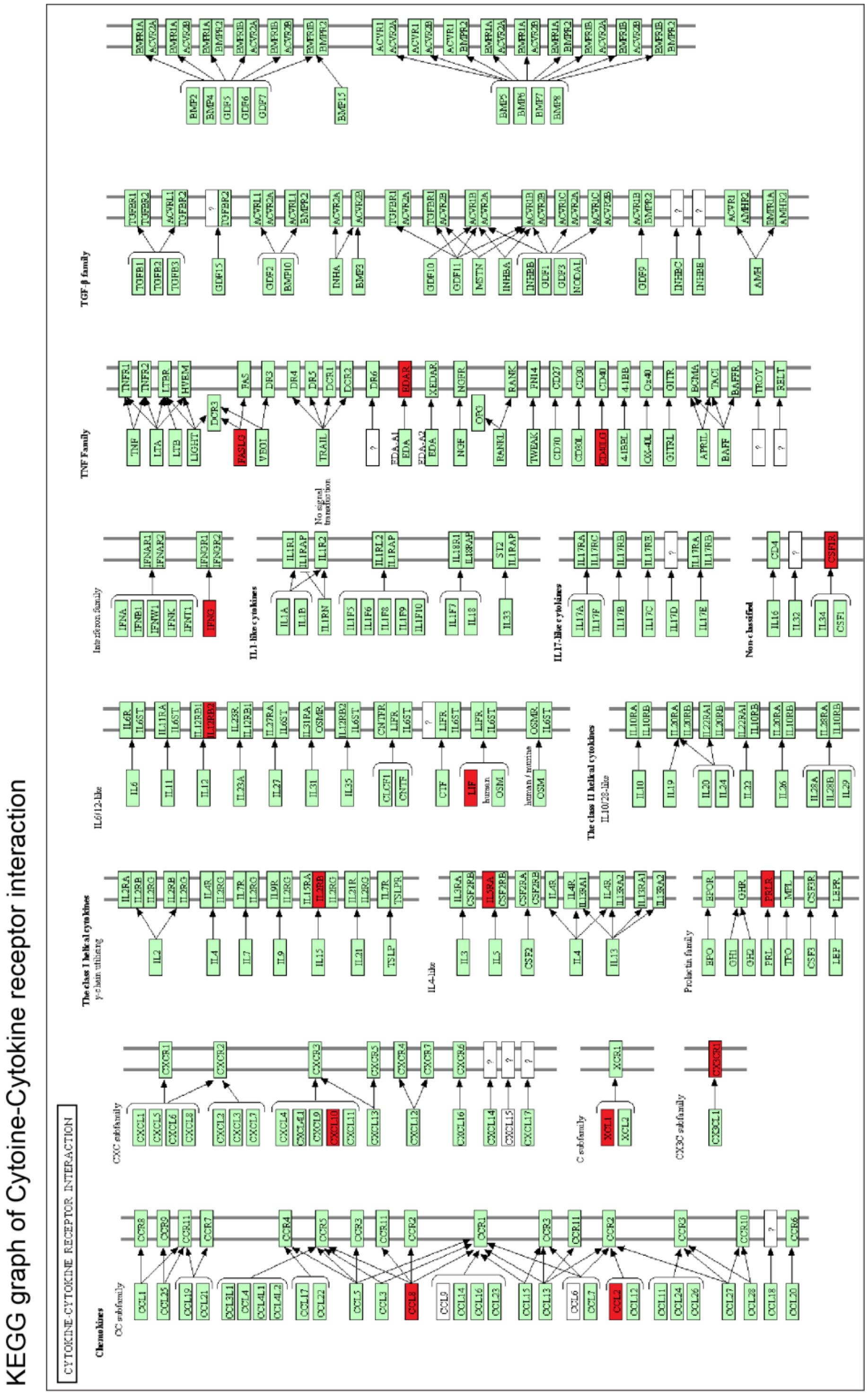


**B**

**Figure 4 - figure supplement 1.** KEGG graphs show genes differentially expressed over time of the Th1 and Th2 cell differentiation pathway **(A)** and the Cytokine-cytokine receptor interaction pathway. **(B)** Red boxes depict up-regulated genes in mild COVID-19 patients during the acute phase of the disease, whereas green boxes depict genes without significant differential gene expression within the KEGG pathways.

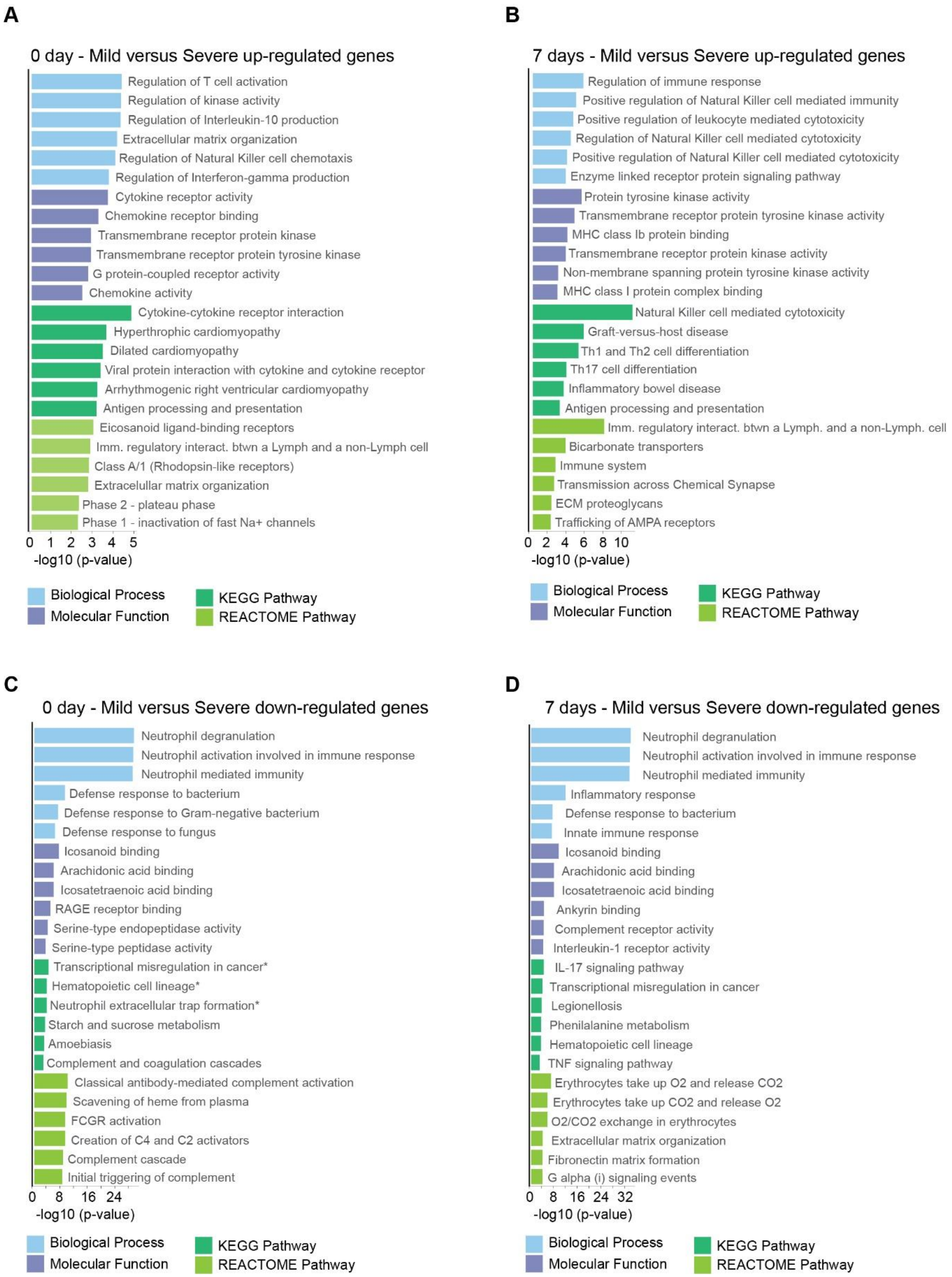


**Figure 4 - figure supplement 2.** Gene Ontology (GO) and KEGG and REACTOME pathways analyses of differentially expressed genes (DEGs) found in the pairwise comparison between D0 and D7 of COVID-19 infection. Bar graphs showing the enrichment of GO biological processes (in blue color),

GO molecular functions (in purple color), KEGG pathways (in dark green color), and Reactome pathways (in light green color) between mild and severe COVID-19 patients. Enrichment results are sorted by -log10(*p*-value) (higher on top) with a cut-off for DEGs ≥ 4 fold-change considering the up-regulated genes at D0 **(A)**, up-regulated genes at D7 **(B)**, down-regulated genes at D0 **(C)**, and down-regulated genes at D7 **(D)**.

**A** Day 0 - KEGG graph of Natural Killer cell mediated cytotoxicity pathway

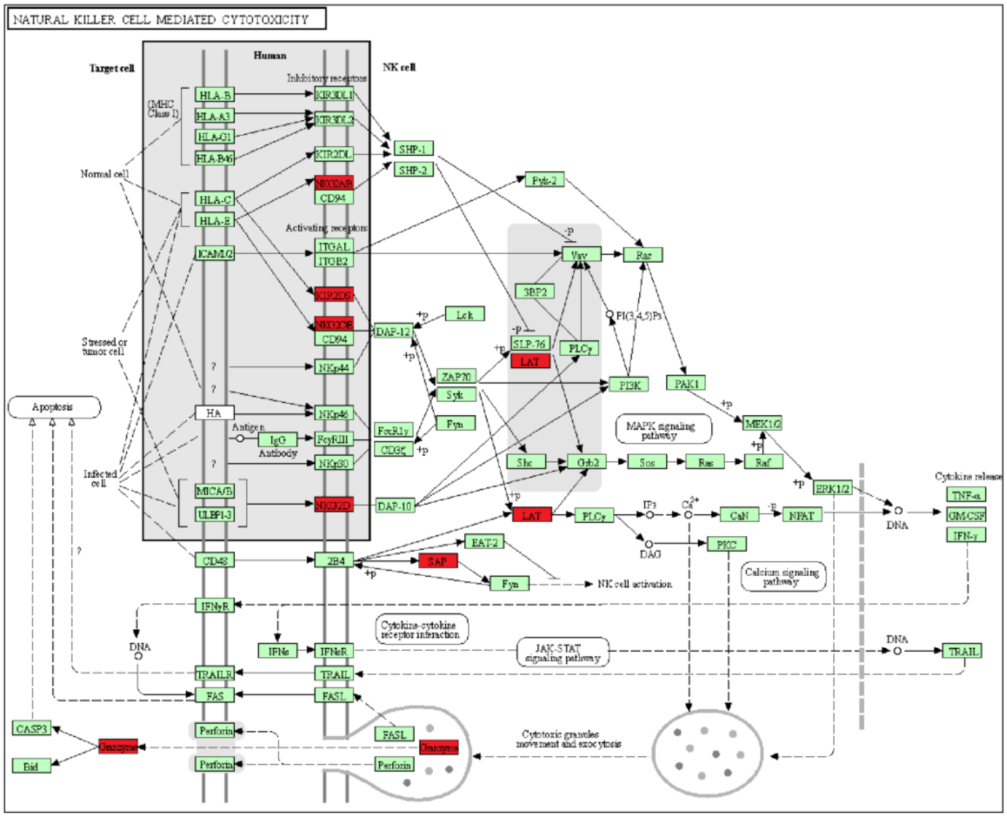


**B** Day 7 - KEGG graph of Natural Killer cell mediated cytotoxicity pathway

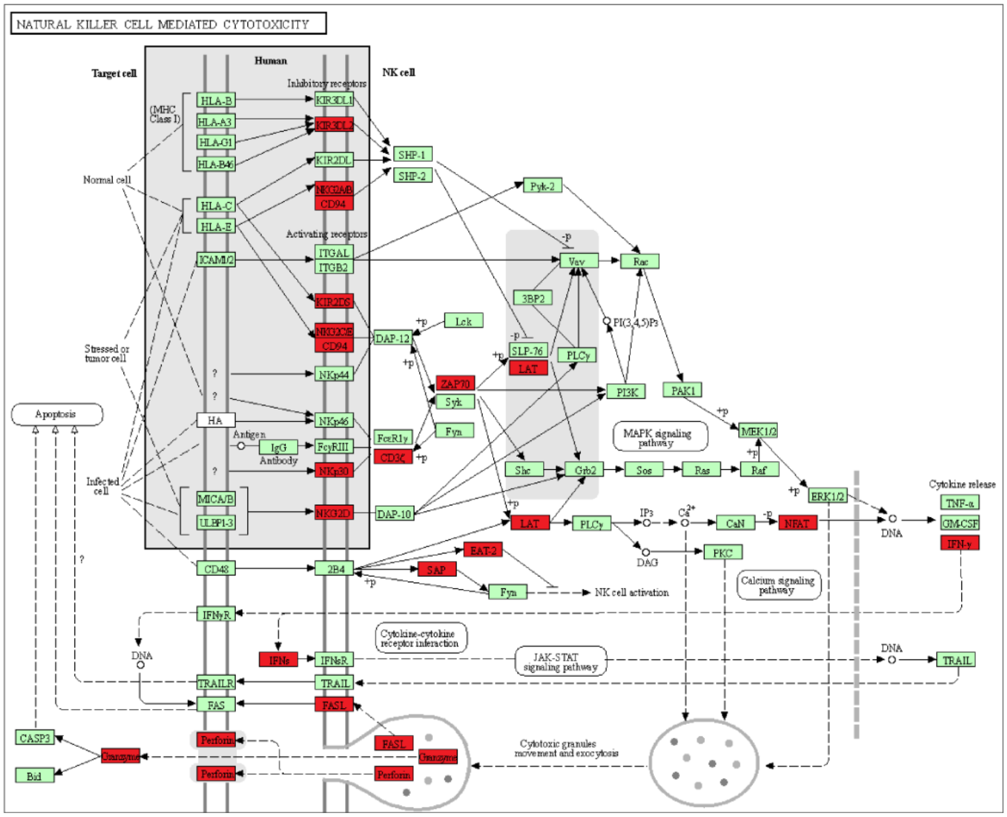

**C** Day 0 - KEGG graph of Th1 and Th2 cell differentiation

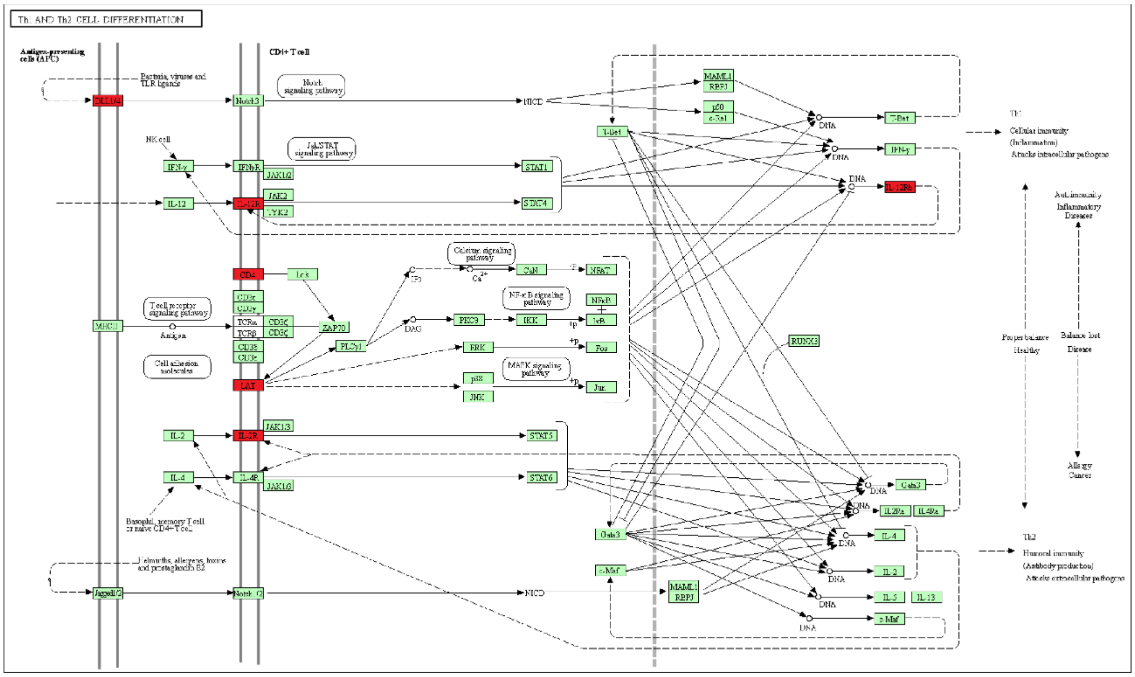


**D** Day 7 - KEGG graph of Th1 and Th2 cell differentiation

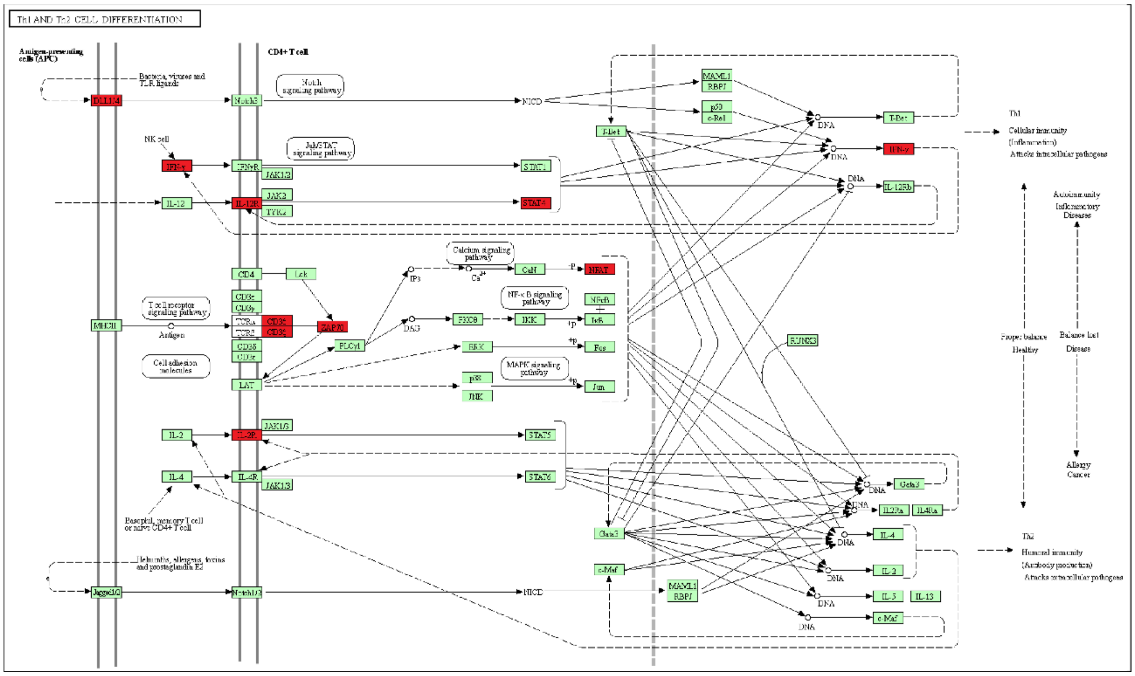

**E** Day 0 - KEGG graph of Cytokine-cytokine receptor interaction

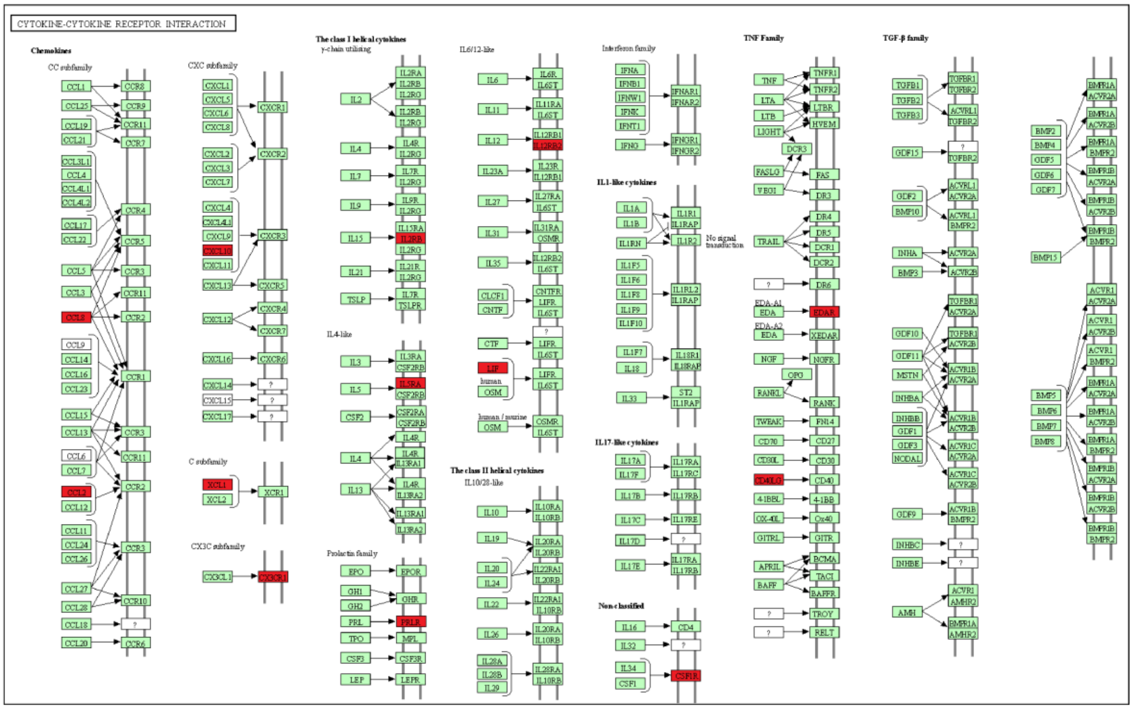


**Figure 4 - figure supplement 3.** KEGG graph show genes with differential expression found in the pairwise comparison (D0 vs D7) from Natural Killer cell-mediated cytotoxicity pathway for up-regulated genes in mild COVID-19 patients at D0 **(A)** and D7 **(B)**, from Th1 and Th2 cell differentiation pathway at D0 **(C)** and D7 **(D)**, and from cytokine-cytokine receptor interaction pathway at D0 **(E)**. Red boxes depict up-regulated genes, whereas green boxes depict genes without significant differential gene expression within each KEGG pathway.

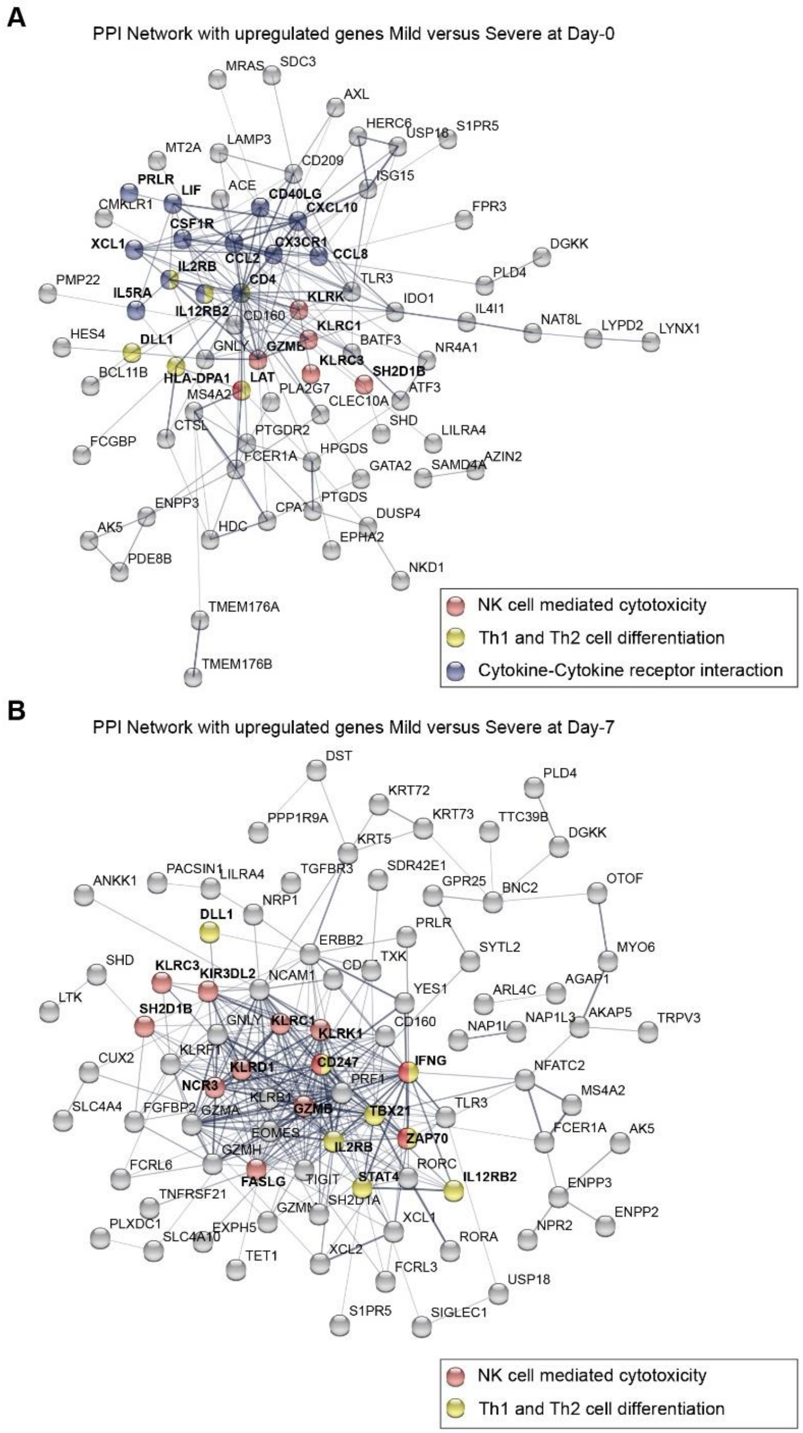


**Figure 4 - figure supplement 4.** Protein-protein interaction (PPI) network graphs show the up-regulated genes found in the pairwise comparison (D0 and D7) in mild versus severe COVID-19 patients. **(A)** Topological representation of the PPI network of up-regulated genes in mild patients on D0. **(B)** Topological representation of the PPI network of up-regulated genes in mild patients on D7. Some nodes are color-coded to highlight proteins involved in the following pathways: Cytokine-cytokine receptor interaction (blue), NK cell-mediated cytotoxicity (red); and Th1 and Th2 cell differentiation (yellow).

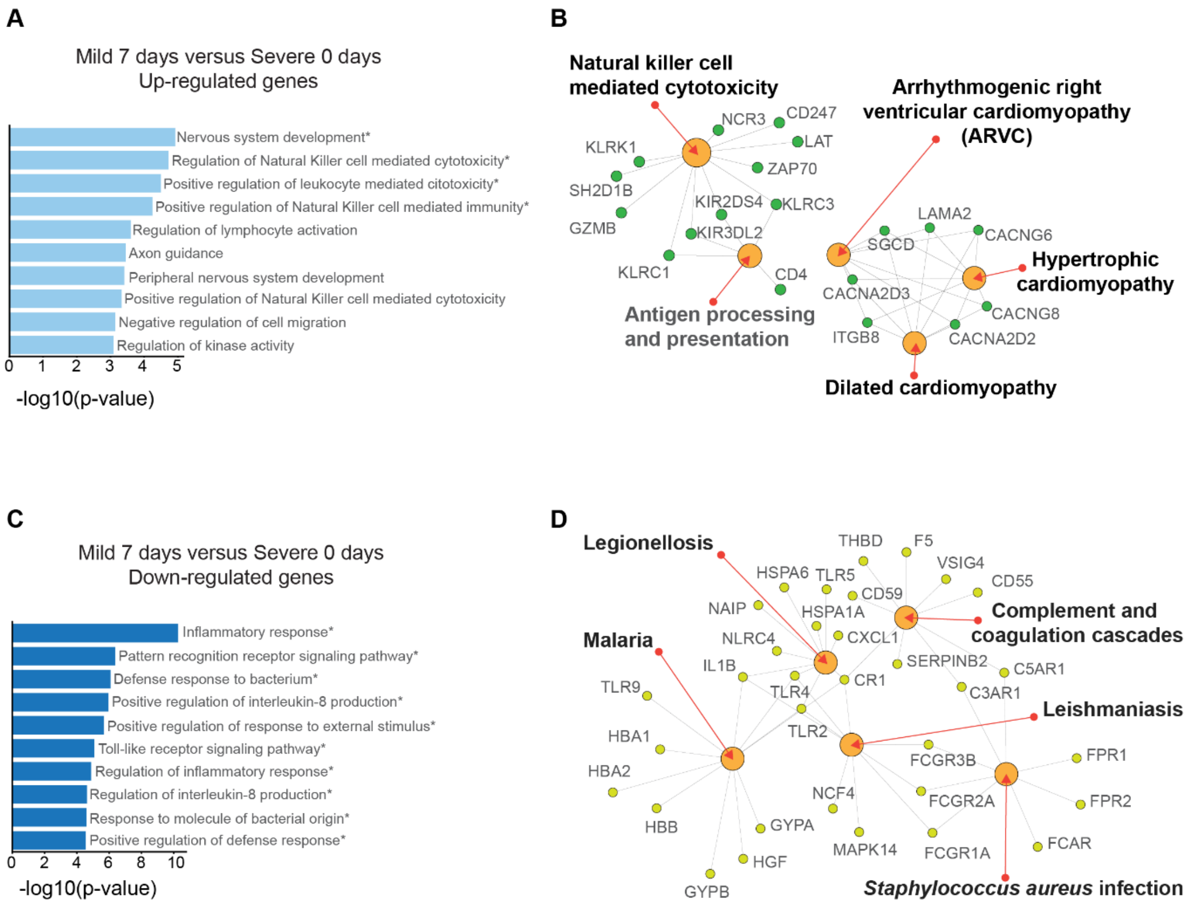


**Figure 4 - figure supplement 5.** Gene ontology and networks of enriched pathways analyses of differentially expressed genes (DEGs) found in the pairwise comparison between D7 of mild patients and D0 of severe patients with COVID-19. Bar graphs show the enrichment of GO terms for biological processes while the networks reveal KEGG enriched pathways for upregulated genes **(A-B)** and downregulated genes **(C-D)** in mild patients, respectively.

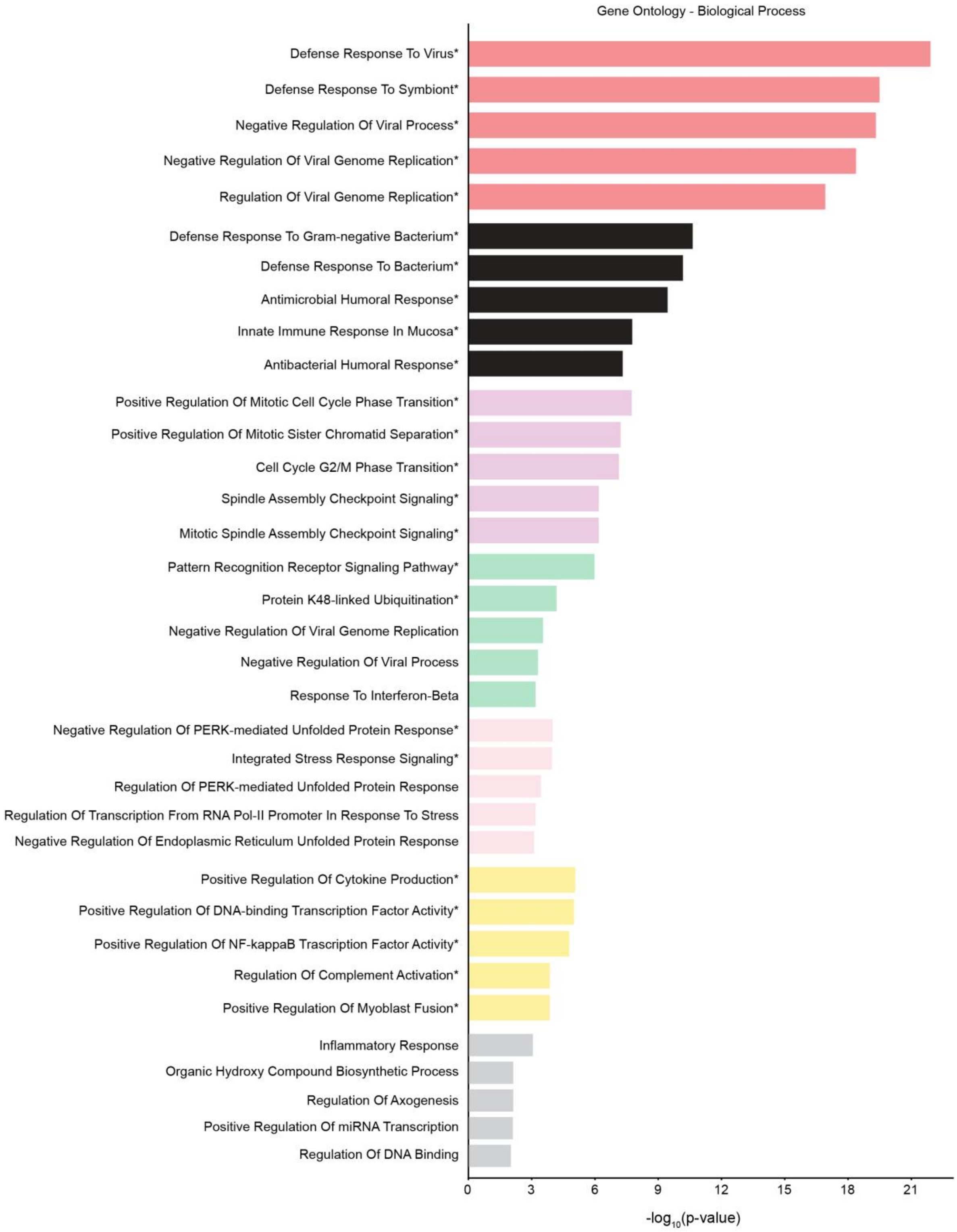


**Figure 5 - figure supplement 1**. Gene ontology analysis of the seven smallest modules of co-expression. The genes from the modules red, black, magenta, green, pink, yellow, and gray, were used to study GO biological process terms. GO terms with statistically significant q-values are marked with an asterisk.

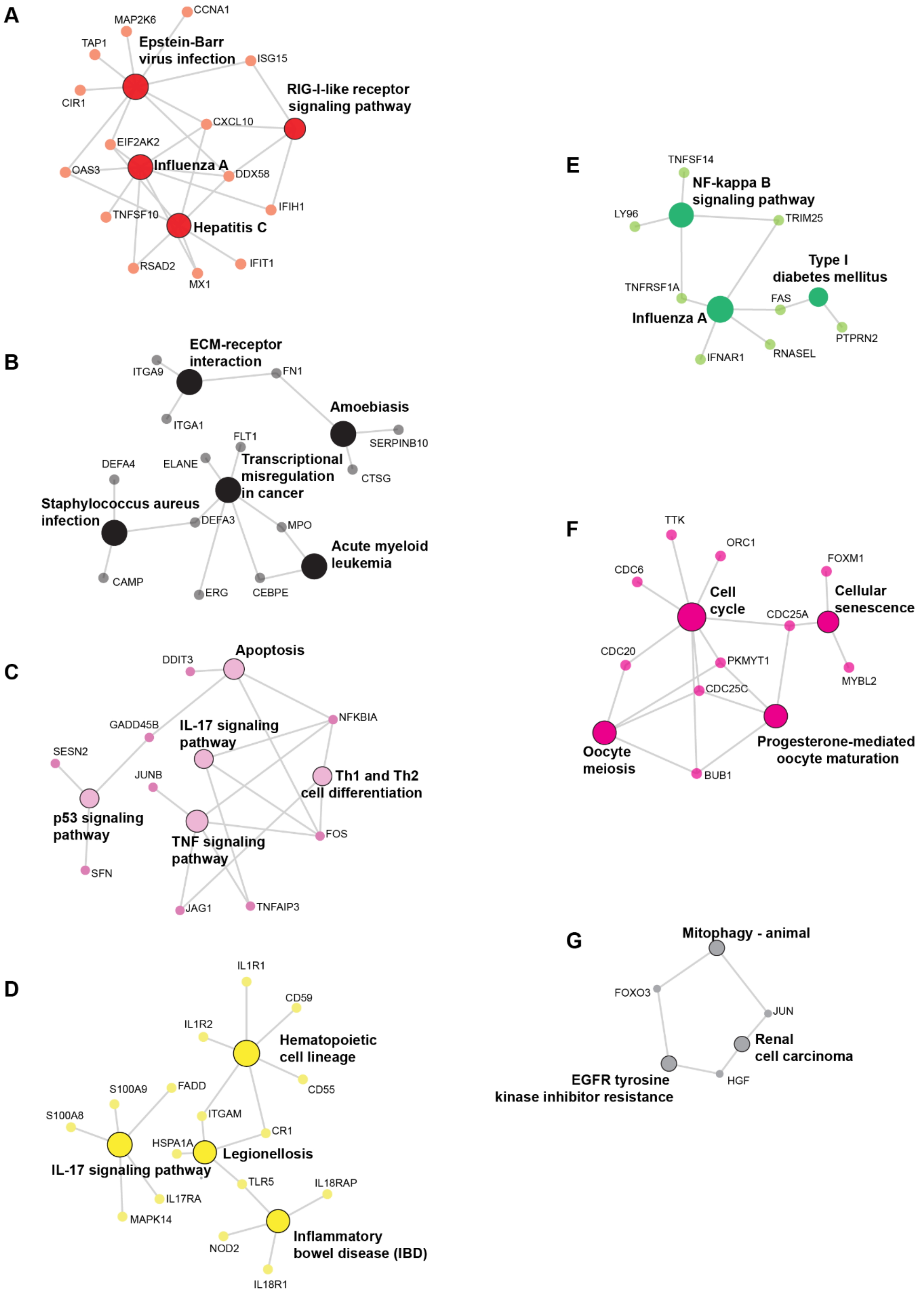
A
Epstein-Barr virus infection
RIG-I-like receptor signaling pathway
Influenza A
Hepatitis C
CCNA1
MAP2K6
TAP1
ISG15
CIR1
CXCL10
EIF2AK2
OAS3
DDX58
IFIH1
TNFSF10
RSAD2
MX1
IFIT1
B
ECM-receptor interaction
Amoebiasis
Transcriptional misregulation in cancer
Staphylococcus aureus infection
Acute myeloid leukemia
ITGA9
FN1
ITGA1
SERPINB10
FLT1
DEFA4
ELANE
CTSG
DEFA3
MPO
CAMP
ERG
CEBPE
C
Apoptosis
IL-17 signaling pathway
Th1 and Th2 cell differentiation
p53 signaling pathway
TNF signaling pathway
DDIT3
NFKBIA
GADD45B
SESN2
JUNB
FOS
SFN
JAG1
TNFAIP3
D
Hematopoietic cell lineage
IL-17 signaling pathway
Legionellosis
Inflammatory bowel disease (IBD)
IL1R1
CD59
IL1R2
S100A9
FADD
CD55
S100A8
ITGAM
CR1
HSPA1A
TLR5
IL18RAP
IL17RA
MAPK14
NOD2
IL18R1
E
NF-kappa B signaling pathway
Type I diabetes mellitus
Influenza A
TNFSF14
LY96
TRIM25
TNFRSF1A
FAS
PTPRN2
RNASEL
IFNAR1
F
Cell cycle
Cellular senescence
Oocyte meiosis
Progesterone-mediated oocyte maturation
TTK
ORC1
FOXM1
CDC6
CDC25A
CDC20
PKMYT1
MYBL2
CDC25C
BUB1
G
Mitophagy - animal
Renal cell carcinoma
EGFR tyrosine kinase inhibitor resistance
FOXO3
JUN
HGF

**Figure 5 - figure supplement 1.** Network for genes of enriched pathways from the seven smallest modules of co-expression. Genes from modules red **(A)**, black **(B)**, pink **(C)**, yellow **(D)**, green **(E)**, magenta **(F)**, and gray **(G)** were used for analysis and the networks show the most enriched pathways.

**Figure 3 - table supplement 1**. Enriched pathways from genes upregulated in mild COVID-19 patients by longitudinal analysis.

**MILD - DAY 0**

| | |
|---|---|
| Natural killer cell mediated cytotoxicity | KLRC3, SH2D1A, PRF1, GZMB, FASLG, KIR3DL2, ZAP70, NCR3, IFNG, KLRD1, KLRC1, CD247, LAT |
| Graft-versus-host disease | IFNG, PRF1, GZMB, FASLG, KLRD1, KLRC1, KIR3DL2 |
| Th1 and Th2 cell differentiation | ZAP70, CD4, IFNG, IL2RB, STAT4, CD247, DLL1, LAT, IL12RB2 |
| Cytokine-cytokine receptor interaction | CX3CR1, CSF1R, CSF3R, LIF, TNFRSF10C, IL5RA, FASLG, PRLR, EDAR, CXCL10, CD4, CD40LG, IL18RAP, CCL8, CXCR1, IFNG, IL2RB, XCL1, CCL2, IL12RB2 |
| Hypertrophic cardiomyopathy | MYBPC3, CACNG6, CACNG8, SGCD, ACE, LAMA2, CACNA2D3, ITGB8, CACNA2D2 |

**MILD - DAY 7**

| | |
|---|---|
| Natural killer cell mediated cytotoxicity | KLRC3, SH2D1A, PRF1, GZMB, FASLG, KIR3DL2, ZAP70, NCR3, IFNG, KLRD1, KLRC1, CD247, LAT |
| Graft-versus-host disease | IFNG, PRF1, GZMB, FASLG, KLRD1, KLRC1, KIR3DL2 |
| Th1 and Th2 cell differentiation | ZAP70, CD4, IFNG, IL2RB, STAT4, CD247, DLL1, LAT, IL12RB2 |
| Hypertrophic cardiomyopathy | CACNG6, CACNG8, SGCD, ACE, LAMA2, CACNA2D3, ITGB8, CACNA2D2, ITGA9 |
| Dilated cardiomyopathy | CACNG6, CACNG8, SGCD, LAMA2, CACNA2D3, ITGB8, CACNA2D2, ADRB1, ITGA9 |

**MILD - DAY 28**

| | |
|---|---|
| Cytokine-cytokine receptor interaction | EDAR, BMP2, CSF3R, IL18RAP, IFNG |
| Pathways in cancer | BMP2, CSF3R, HEY1, IFNG, FLT4, LPAR2, KIF7 |
| Malaria | IFNG, KLRB1 |
| Basal cell carcinoma | BMP2, KIF7 |
| Inflammatory bowel disease | IL18RAP, IFNG |

79 **Figure 3 - table supplement 2**. Enriched pathways from genes upregulated in severe COVID-19 patients by longitudinal analysis.

**SEVERE - DAY 0**

| | |
|---|---|
| Neutrophil extracellular trap formation | H2BC7, CR1, H2BC6, AQP9, NCF4, C5AR1, FPR1, FPR2, MAPK14, MPO, TLR8, PADI4, H4C4, FCGR1A, ELANE, TLR2 |
| Hematopoietic cell lineage | GYPA, CSF3R, IL4R, CR1, MME, IL1R1, FLT3, IL1R2, FCGR1A, CD55 |
| Viral protein interaction with cytokine and cytokine receptor | IL18RAP, TNFSF14, CXCR1, TNFRSF10C, CXCL1, CXCL2, IL18R1 |
| Amoebiasis | SERPINB10, IL1R1, ARG1, IL1R2, FN1, CXCL1, CXCL2, TLR2 |
| Complement and coagulation cascades | THBD, CR1, SERPINB2, PLAU, C5AR1, VSIG4, CD55, F5 |

**SEVERE - DAY 7**

| | |
|---|---|
| Transcriptional misregulation in cancer | CEBPB, FLT1, BCL2A1, GADD45A, FLT3, CEBPE, IL1R2, DEFA4, ZBTB16, DEFA3, MPO, MMP9, GADD45G, BCL6, PLAU, PPARG, ERG, FCGR1A, ELANE |
| Neutrophil extracellular trap formation | CR1, AQP9, NCF4, C5AR1, FPR1, FPR2, AZU1, MAPK14, MPO, TLR8, CTSG, PADI4, FCGR1A, ELANE, CAMP, TLR2 |
| IL-17 signaling pathway | FOSL1, CXCL6, CEBPB, LCN2, TNFAIP3, CXCL1, MAPK14, CXCL2, MMP9, S100A9, S100A8 |
| TNF signaling pathway | SOCS3, CXCL6, CEBPB, TNFAIP3, CXCL1, MAPK14, CXCL2, MMP9, IL18R1, CREB5 |
| Hematopoietic cell lineage | GYPA, CSF3R, IL4R, CR1, MME, IL1R1, FLT3, IL1R2, CD24, FCGR1A, CD55 |

**SEVERE - DAY 28**

| | |
|---|---|
| Malaria | CR1, TLR9, SDC1, THBS1, TLR2 |
| Legionellosis | CR1, TLR5, CXCL2, TLR2, HSPA1A |
| Neutrophil extracellular trap formation | CR1, C5AR1, FPR1, TLR8, FPR2, FCGR1A, MAPK14, TLR2 |
| Toll-like receptor signaling pathway | CXCL10, TLR9, TLR8, TLR5, MAPK14, TLR2 |
| Complement and coagulation cascades | THBD, CR1, SERPINB2, C5AR1, CD55 |
